# Multi-finger Receptive Field Properties in Primary Somatosensory Cortex: A Revised Account of the Spatio-Temporal Integration Functions of Area 3b

**DOI:** 10.1101/2022.03.21.485210

**Authors:** Natalie K. Trzcinski, S.S. Hsiao, Charles E. Connor, Manuel Gomez-Ramirez

## Abstract

The leading view in the somatosensory system indicates that area 3b serves as a cortical relay site that encodes cutaneous (tactile) features limited to individual digits. Our recent work argues against this model by showing that cells in area 3b integrate information from cutaneous and proprioceptive modalities. Here, we further test this model, by studying the multi-finger neural integration properties of area 3b. In contrast to the prevailing view, we found that most cells in area 3b have a receptive field (RF) that extends to multiple digits. Responses to tactile stimulation emerged earlier in cells with a multi-digit (MD) vs. single-digit (SD) RF. We also found that the RF size of MD cells (the number of responsive digits) increased across time, and the orientation preference across digits was highly correlated. Taken together, these data provide strong evidence that area 3b plays a larger role in generating neural representations of tactile objects, as opposed to just being a ‘feature detector’ relay site.

## Introduction

Object sensing and manipulation (i.e., haptics) is largely mediated by brain mechanisms that integrate tactile information across fingers. Haptics commences in the periphery with specialized cells that transduce objects’ physical features into neural signals that are relayed and combined in the brain (Hsiao, 2008; Hsiao and Gomez-Ramirez, 2013; Mountcastle, 2005). As information is relayed to the brain, the receptive field (RF) of cells increases and becomes more elaborate. Peripheral cells with the highest spatial acuity (i.e., Merkel cells) are confined to small skin areas (∼2-8 mm), whereas RFs in area 3b have been thought to be a single finger (Iwamura et al., 1983, 1994; Pons et al., 1987; Sur, 1980). In cortical areas downstream of area 3b, RFs tend to encompass multiple fingers (e.g., area 1) or the whole hand (e.g., area 2 and secondary somatosensory cortex; *SII*), and have complex inhibitory and excitatory spatio-temporal properties (DiCarlo et al., 1998; Fitzgerald et al., 2004; Sripati, 2006; Thakur et al., 2006). Cells in area 3b, and to a lesser extent area 1, are thought to operate as ‘local feature detectors’, with RFs akin to those of simple cells in primary visual (V1) cortex (Bensmaia et al., 2008). These observations have led to the view that tactile object perception emerges across cortical stages, likely in area 2 or beyond, with area 3b serving as a cortical relay site that encodes tactile features confined to individual digits (Hsiao, 2008).

Our understanding of area 3b cells’ RF size is largely derived from anesthetized and awake-behaving studies in non-human primates that primarily analyzed responses to stimulation on a single finger (Ageranioti-Bélanger and Chapman, 1992; Bensmaia et al., 2008; Costanzo and Gardner, 1980; DiCarlo et al., 1998; Gardner et al., 1999; Pei et al., 2009, 2010; Pons et al., 1987; Sripati, 2006; Sur, 1980; Sur et al., 1984, 1985; Warren et al.). Although much was learned from these studies, anesthesia can have a significant impact on neurons’ activation strength (Hildebrandt et al., 2017; Krom et al., 2020; Sorrenti et al., 2021), potentially undermining weak, but reliable, responses to stimulation of different fingers. Further, the vast majority of area 3b studies in awake-behaving animals are also limited because researchers failed to properly assay responses to tactile stimulation in every finger. In these awake-behaving studies (as well as anesthetized studies), researchers typically localized the finger with the maximum response (i.e., the preferred finger) using unreliable hand held probes, and then used motorized devices to systematically stimulate the preferred finger only (see e.g.,(Ageranioti-Bélanger and Chapman, 1992; Bensmaia et al., 2008; DiCarlo et al., 1998; Pei et al., 2009, 2010, 2011; Sripati, 2006; Tabot et al., 2013)). The concern with this approach is that we may have an incomplete, and likely incorrect, representation of the cross-finger neural integration properties of area 3b cells. Indeed, using automatized stimulators, we showed that area 3b cells integrate inputs from proprioceptive and cutaneous modalities (Kim et al., 2015), a finding that runs counter to the modality specificity model of the somatosensory system (Mountcastle, 2005; Sur et al., 1981, 1984). Thus, there is a need to comprehensively characterize area 3b’s RFs using automated and systematic approaches.

To the best of our knowledge, there has only been one study that recorded activity in area 3b while stimulating multiple locations on the hand using reliable motorized tools (Reed et al., 2010a). Co-stimulation of the preferred skin region and a neighboring site modulated responses relative to stimulation on the preferred skin region alone, suggesting that area 3b cells have multi-digit (MD) non-classical RFs. However, this study failed to systematically test responses across all digits, and collapsed across single and multi-unit spiking data. Many cells reported in (Reed et al., 2010a) were observed in boundary locations of the digit representations. Therefore, it is unclear whether cells classified with a MD RF truly emanate from an individual cell or multiple cells. Further, it is still unknown what mechanisms give rise to these MD RFs, and how information across fingers is represented by MD cells.

To address these questions, we performed single-unit recordings in the finger representation of area 3b while systematically presenting oriented bars to different fingers in awake-behaving monkeys. Contrary to the prevailing model, we found that the majority cells in area 3b have RFs that extend to multiple digits. MD RFs evolved as a function of time, suggesting that they are mediated by feedback mechanisms. We also found that MD cells tuned to oriented bars have an orientation preference across fingers that differ between 22.5° and 67.5°, an ideal range for encoding curved objects. Taken together, these data provide compelling evidence that area 3b integrates neural information across fingers, indicating a more integral role in mediating object representations than previously thought.

## Results

Two animals (*Macaca mulatta*) sat comfortably on a custom-made chair with their hands held supinated (**Figure 1**). We stimulated digits 2, 3, 4, or 5 while monkeys performed an unrelated visual contrast discrimination task. Tactile stimuli consisted of oriented bars (0° – 157.5°, steps of 22.5°, indented 1mm into the skin for 500ms duration). Single unit activity was recorded using a custom-built electrode microdrive with four linearly-space electrodes from four hemispheres of the distal finger representation of area 3b (see Methods section). We used similar functional mapping procedures as in previous studies (Bensmaia et al., 2008; DiCarlo et al., 1998; Pei et al., 2009, 2010, 2011; Thakur et al., 2012) to ensure single unit recordings were made in area 3b. Analyses were conducted in cells recorded >2mm below the surface of the brain to avoid analyzing cells in area 1. The anatomical organization of areas 1 and 3b in rhesus monkeys is highlighted by the inverse topography between the distal, medial and proximal finger pad representation in areas 1 and 3b (Mountcastle, 2005). Recording from the proximal pad representation in area 3b may lead to the inclusion of area 1 cells with a RF over the proximal pad. As such, we only selected cells in area 3b with a RF over the distal finger pad.

**Figure 1:**
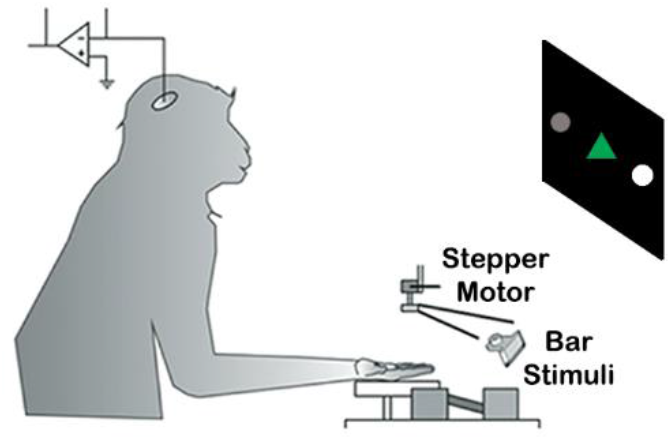
Experimental Setup. Animals set in a custom-made chair with their hand supinated and secured. Bars with different orientations were systematically delivered to digits 2 through 5 while animals were engaged in a visual contrast discrimination task. Animals were trained to fixate on a green triangle, and saccade to the flanking circle with highest brightness. After animals were fully trained on the visual task we recorded single-unit activity from the distal representation of area 3b.

### Area 3b cells have RFs that cover multiple digits

The majority of area 3b cells (∼57%) responded to stimulation on more than one digit, a finding that runs counter to the leading model of RF size in the somatosensory system (Pons et al., 1987; Sur, 1980; Sur et al., 1980). **Figure 2A** shows examples of neurons with a RF over one (top row), two (second row), three (third row), or four (bottom row) digit(s). The insets in **Figure 2A** show averaged action potential waveforms recorded to each stimulated digit. **Supplementary Figure 1** shows additional examples of area 3b neurons with RFs over 1, 2, 3 or 4 digits. **Figure 2B** shows the RF size distribution across the population. As expected, we observed a reduction in the number of cells with MD RFs, with ∼43% of cells having a RF confined to a single finger, and ∼10% of cells having a RF size that covered four digits. This pattern of effects was observed in each of the hemispheres recorded in both monkeys (**Supplementary Figure 2**). A Kruskal-Wallis test showed that the firing rate of the preferred digit was significantly greater in neurons with larger RF size (H(3,473) = 83.19, p < 0.001; **Figure 2C**). A regression test showed a systematic relationship between firing rate and RF size (R^2^ = .15, p < 0.001). Similar findings were observed during the baseline firing rate (H(3,473) = 19.56, p < 0.001; **Supplementary Figure 3**), and the firing rates to the 2^nd^ preferred digit (H(2,269) = 46.03, p < 0.001; **Supplementary Figure 4**). The preferred digit location was uniformly distributed across all fingers, indicating that we equally sampled neurons from all finger representations of area 3b (**Supplementary Figure 5A**, X^2^(3,473) = 0.57, *p > 0.05*). Amongst cells with a MD RF, we found that the second most responsive digit was predominantly adjacent to the preferred digit (**Supplementary Figure 5B**, X^2^(5,269) = 160.54, *p < 0.001*).

**Figure 2:**
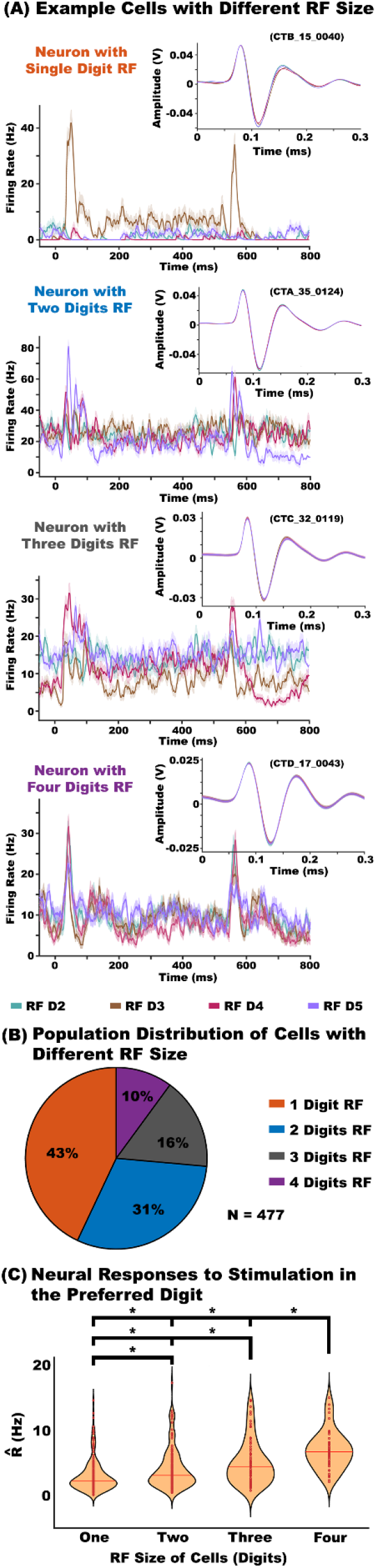
Majority of Cells in Area 3b Have RFs Spanning Multiple Digits. **(A)** Examples neurons a RF spanning one (top row), two (second row), three (third row), or four (bottom row) digits. The insets in each graph show action potential waveforms recorded to each stimulated digit. **(B)** RF size distribution across the recorded population. Most cells had a RF covering 2 or more digits (∼57% of cells). However, cells with a SD had the largest incidence (∼43%), whereas cells with a 4 digits RF had the lowest incidence (∼10%). **(C)** Firing rates to stimulation on the preferred digit increased as a function of cells’ RF size. The graph shows violin plots of the normalized firing rate (R̂) distribution for each RF size condition. Each dot represents a cell’s mean R̂. P value < 0.01 = *.

The firing rate response in MD cells was not uniform across digits, indicating that MD cells have a digit that elicit a preferential response (i.e., a preferred digit). **Figure 3** shows the cumulative distribution of the RF Fano Factor (RF_FF_, see Methods) across all cells, which measures a neuron’s response similarity across digits. An RF_FF_ = 1 indicates a MD cell with Poisson distributed responses across digits. An RF_FF_ < 1 or RF_FF_ > 1 indicates a MD cell with similar responses across digits or unequal responses across digits (i.e., a MD cell with a preferred digit), respectively. A linear regression showed that the RF_FF_ significantly increased as a function of RF size (R^2^ = 0.03; p < 0.01). The inset of **Figure 3** shows the median RF_FF_ across RF digit size. Increases in RF_FF_ are likely due to increased response variability across fingers (i.e., cells having non-homogeneous response functions across responsive digits) since the spiking of cells (i.e., R̂) increased as a function of RF size (**Figure 2C)**.

**Figure 3:**
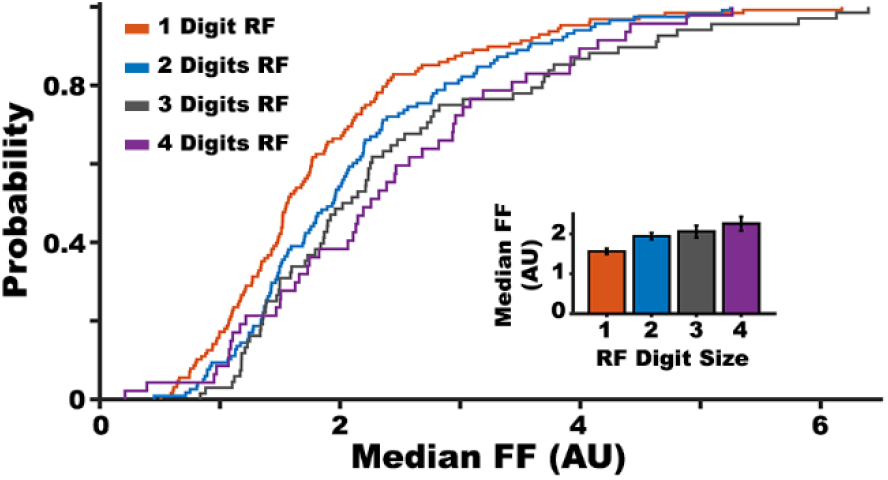
Neural Responses Across are not Uniformly Distributed. Cumulative distribution RF Fano Factor (RF_FF_), which measures a neuron’s response similarity across digits. The data shows a systematic shift in the cumulative distribution across RF size, with four digit RF cells having the largest RF_FF_. The inset shows the median RF_FF_ across RF digit size. These indicate that MD cells have a particular digit eliciting a preferential response.

### MD RF cells respond faster and have higher spike timing precision

Neural response to the preferred digit of MD cells emerge earlier as compared to those in cells with a SD RF (X^2^(1,476) = 24.65, *p < 0.0001*). The upper panels in **Figure 4A** show examples of neural responses to stimulation on the preferred digit in cells with a SD (red trace, left y-axes) and MD (blue trace, right y-axes) RF. The lower panel in **Figure 4A** shows the cumulative distribution of onset time for the preferred digit on SD (red traces) and MD (blue traces) cells with excited (solid trace) and inhibited (dashed trace) initial responses, relative to baseline. As the figure shows, the earliest responses were primarily inhibited (X^2^(1,476) = 65.24, *p < 0.0001*). Importantly, we only considered cells that had an initial response within 300ms following stimulus onset. Somatosensory cortex has cells that respond only to the offset of the stimulus (Gomez-Ramirez et al., 2014; Pei et al., 2009; Reed et al., 2010b, 2010a), thus including these cells in this analyses would introduce biases to the distribution.

**Figure 4:**
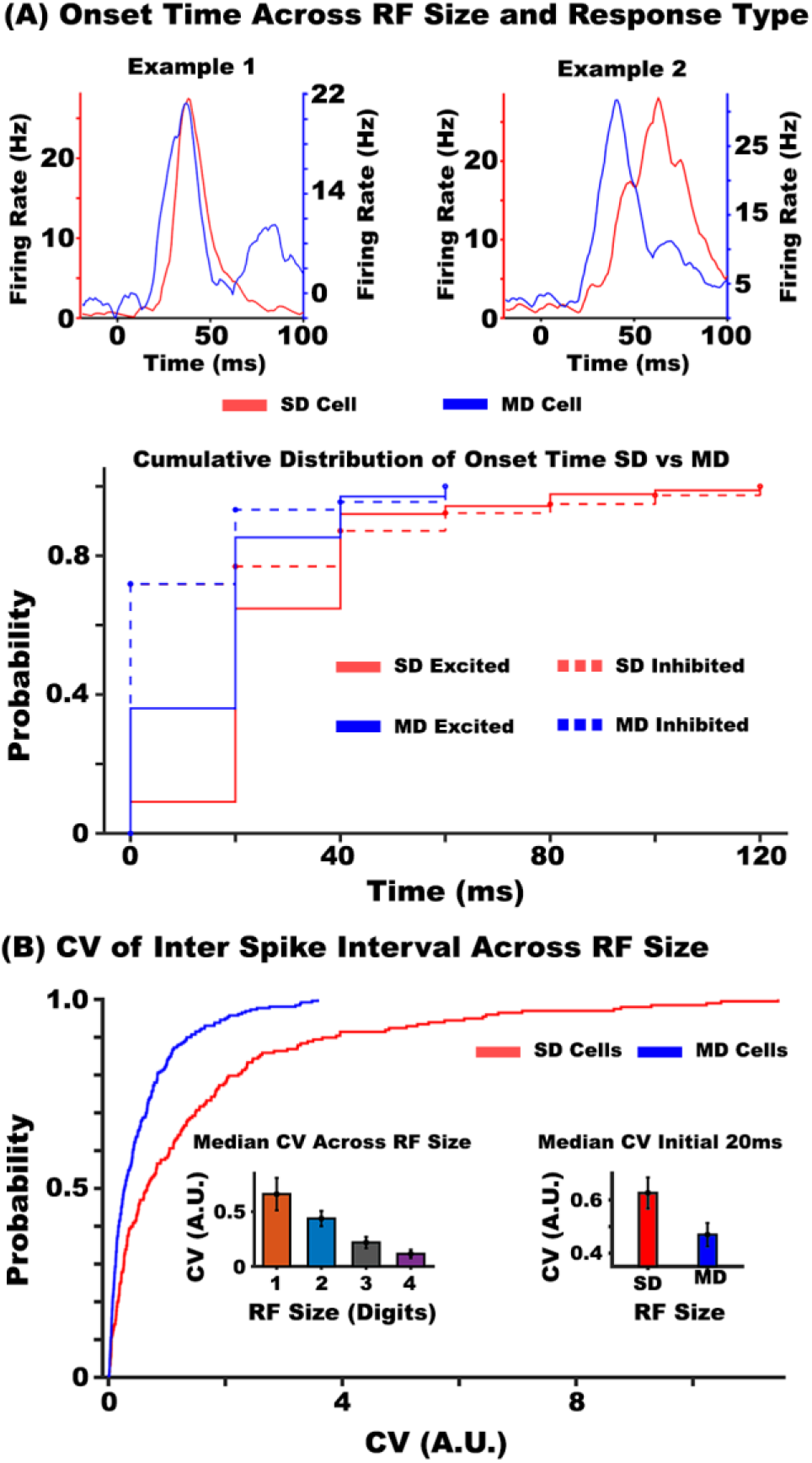
Neural Responses Onset Faster and are More Precise in MD Cells. **(A)** The top graph shows the instantaneous firing rate on the preferred digit of four neurons with excited responses (two in the left panel, two in the right panel) with a SD RF (red trace) and a MD RF (blue trace). Note that the firing rate axes for SD and MD are plotted on the left and right axes, respectively. The lower panel shows the cumulative distribution of onset time for the preferred digit on SD (red traces) and MD (blue traces) cells with excited (solid trace) and inhibited (dashed trace) initial responses. The earliest activations were largely from cells with a response below baseline (i.e., inhibited). **(B)** Cumulative distribution of the Coefficient of Variation (CV) of the inter spike interval (ISI). The CV of the ISI was lower (i.e., more regular) for MD vs. SD cells. The left shows the median CV of the ISI decreasing as a function of RF size. The right inset shows the median CV of the ISI in MD (blue bar) and SD (red bar) cells with a significant response during the initial 20ms relative to stimulus onset.

The spike timing, measured as the Coefficient of Variation (CV) of the inter spike interval (ISI)(Shin and Moore, 2019), was more regular in MD vs. SD cells (Z = 5.69; *p < 0.001*). **Figure 4B** shows the cumulative distribution of the CV of the ISI for SD (red trace) and MD (blue trace). The left inset of **Figure 4B** shows that the median CV of the ISI decreased as a function of RF size (X^2^(3,453) = 50.41, *p < 0.0001)*. As shown in the right inset of **Figure 4B,** cells with a significant response during the first 20ms of stimulus onset exhibited more regular spiking if they had a MD vs. SD RF (Z = 2.17; *p < 0.05)*. These data indicate that MD cells have more reliable feedforward inputs from first-order thalamic sources as compared to cells with SD.

### MD RFs in area 3b evolve across time

Responses to non-preferred digit stimulation (i.e., the size of MD RFs) increased as a function of time, suggesting the MD RFs are generated by feedback (or reentrant) activity from downstream cortical sources. The upper panels in **Figure 5** show examples of neural responses to tactile stimulation on each digit of a cell with a RF spanning four digits. The example cells show a systematic delay in response onset across finger stimulation. The lower panel in **Figure 5** shows the cumulative distribution of onset time for each responsive digit across MD cells. Onset response times were systematically delayed across responsive digits (X^2^(3,473) = 98.05, *p < 0.0001*). The inset in the cumulative distribution of **Figure 5** shows the median response onset time for each responsive digit across the population, with the fastest digit response occurring within the first 20ms, the second fastest digit response occurring between 20 and 40ms, and the remaining digit responses occurring after ∼40ms. A regression analysis showed a systematic relationship between response onset across stimulated fingers (R^2^ = 0.18, *p < 0.0001*).

**Figure 5:**
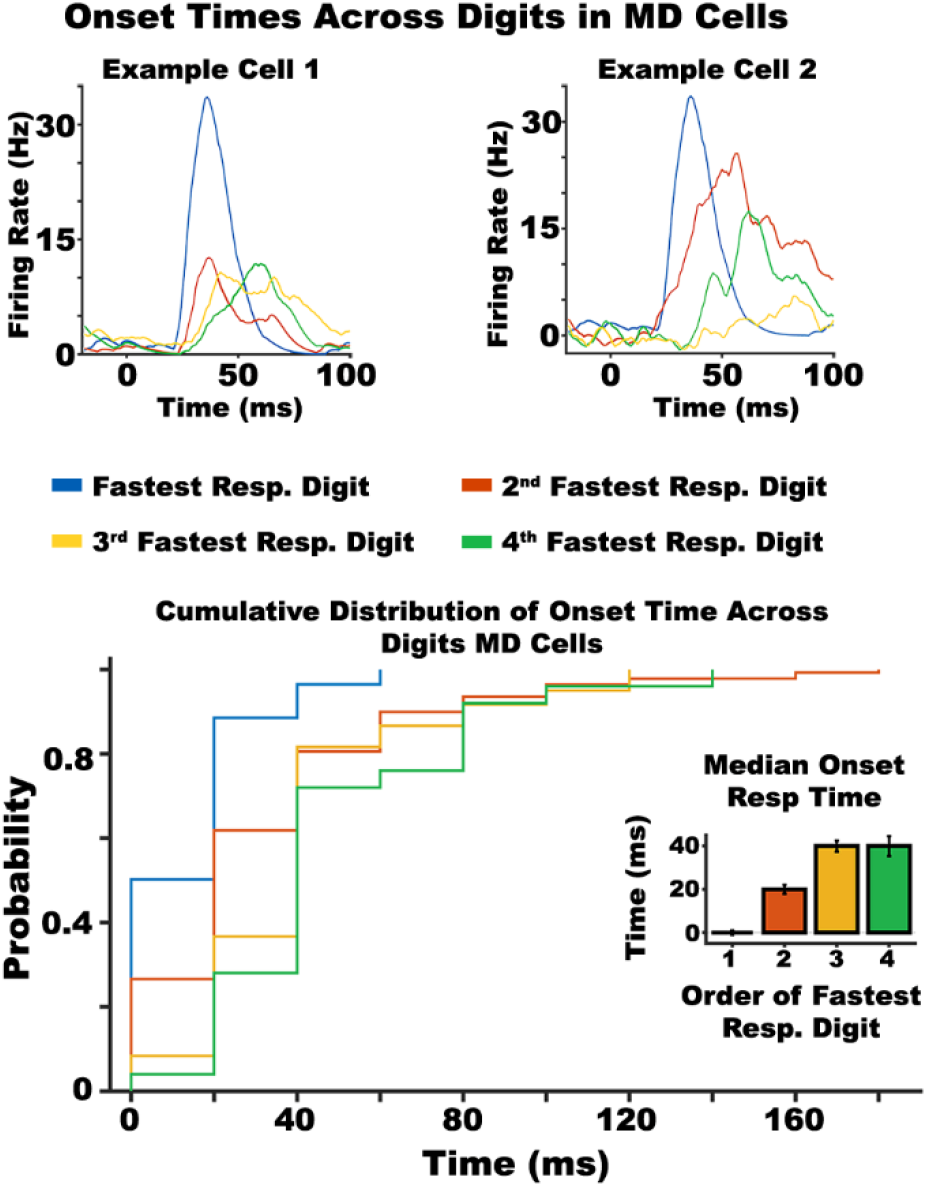
MD RFs Evolve Across Time. The top graphs show examples of neural responses to tactile stimulation on each digit of a cell with a RF spanning four digits. The lower panel shows the cumulative distribution of onset time for each responsive digit across MD cells. The graph shows a systematic shift in response onset times across responsive digits. The inset in the cumulative distribution shows the median response onset time for each responsive digit of MD cells.

### Orientation tuning properties decrease as RF size increases

Orientation tuning was more prevalent in the preferred digit of cells with a SD vs. MD RF (X^2^(3,215) = 124.93, *p < 0.0001*). **Figure 6A** shows examples of orientation tuned cells with a RF in one (upper left panel), two (upper right panel), three (lower left panel), or four (lower right panel) digits. The dashed black line shows the average product of the tuning functions between preferred and non-preferred digits. **Figure 6B** shows the percentage of orientation tuned cells with a one, two, three, or four digit RF. With the exception of cells with a four digits RF (X^2^(3,46) = 20.43, *p < 0.001*), most MD cells with orientation tuning in the preferred digit also show significant orientation tuning in the non-preferred digits. **Figure 6C** shows the orientation index (OI) of cells with a one, two, three, or four digits RF. We observed greater orientation selectivity in SD vs. MD cells (Z = 2.16; *p < 0.05*). However, this difference was largely found between cells with a one vs. three digits RF (approaching significance; Z = 1.89; *p = 0.06*), and one vs. four digits RF (Z = 2.42; *p < 0.05*). We also failed to see significant differences in OI between digits (H(3,458) = 2.83, p > 0.05); **Supplementary Figure 6).**

**Figure 6:**
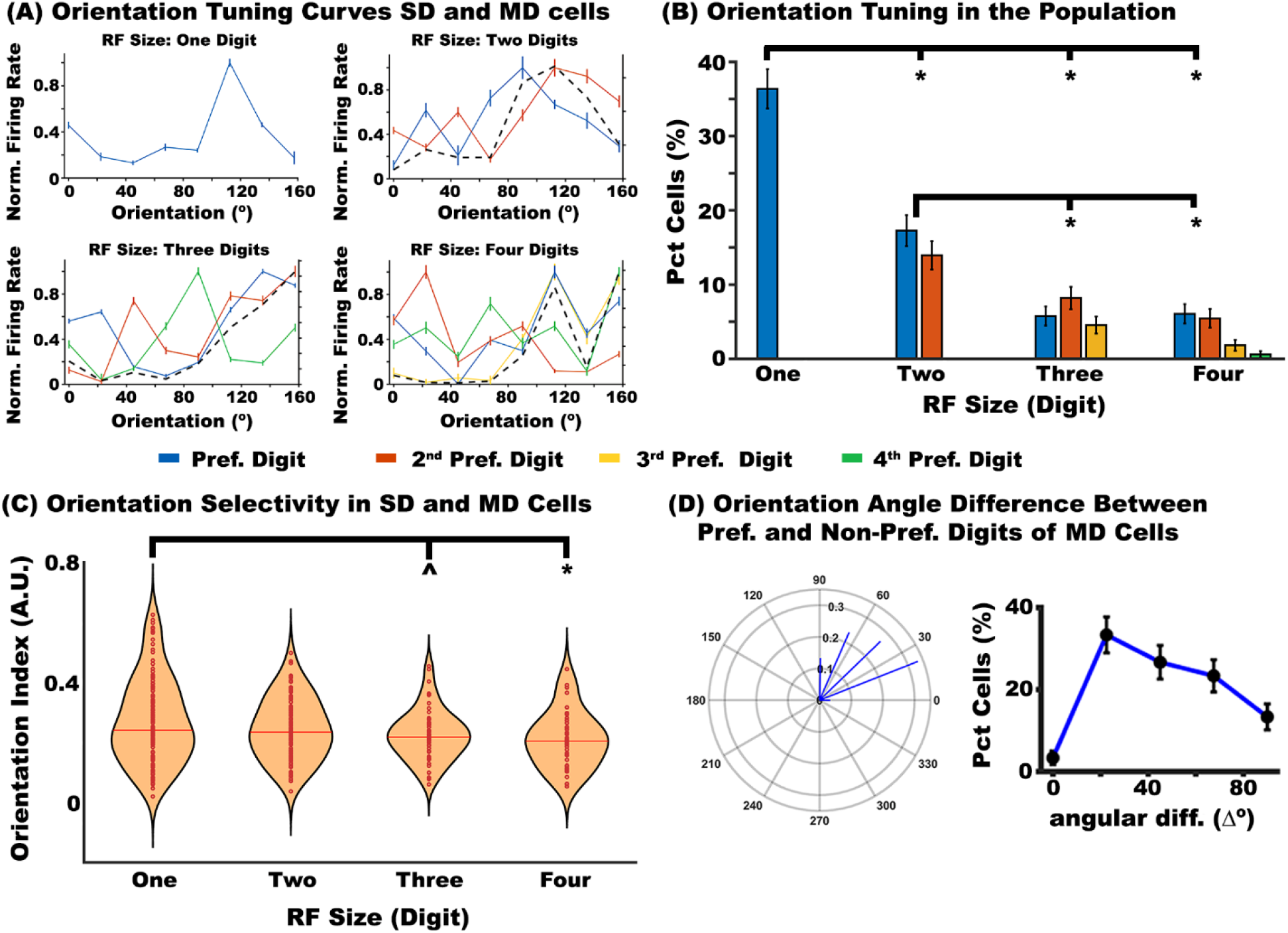
Orientation Tuning Properties of SD and MD Cells. **(A)** Examples orientation tuning curves in cells with one digit (top left graph), two digits (top right graph), three digits (bottom left graph), and four digits RF (bottom right graph). The dashed black line in MD cells shows the average product of the tuning functions between preferred and non-preferred digits. **(B)** The percentage of orientation tuned cells with a one, two, three, or four digit RF. The graph indicates that orientation tuning was more prevalent in the preferred digit of cells with a SD vs. MD RF. **(C)** Orientation index (OI) in the preferred digit of cells with a one, two, three, or four digits RF. The data revealed greater orientation selectivity in SD vs. MD cells. **(D)** The left panel shows a polar plot of the difference in the preferred angle between preferred and non-preferred digits of orientation tuned MD cells. Each line represents the length of the difference vector between preferred and non-preferred digits. The right panel shows the percentage of cells with an angular difference between 0° and 90°. Most MD cells have a preferred angle difference between preferred and non-preferred digits of 22.5°. P value < 0.01 = *; P value = 0.05 = ^.

### Orientation tuning preference of MD cells are correlated across digits

The preferred angle of preferred vs. non-preferred digits of MD cells is shifted between 22.5° and 67.5° (R = 0.43; *p < 0.05;* see black dashed lines in **Figure 6A**). **Figure 6D** (left panel) shows a polar plot of the difference in the preferred angle between preferred and non-preferred digits of orientation tuned MD cells. The right panel of **Figure 6D** shows the percentage of cells with an angular difference between 0° and 90°. Most MD cells have a preferred angle difference between preferred and non-preferred digits of 22.5°, with the least preferred angle difference being 0° (X^2^(4,115) = 12.53, *p < 0.05*).

## Discussion

Our study showed that the majority of cells in area 3b have a RF that spanned multiple digits, a finding that runs counter to the leading view of cells’ RF size properties in the somatosensory system (Iwamura et al., 1983, 1994; Mountcastle, 2005; Pons et al., 1987, 1987; Sur, 1980). We also found that the firing rate response across each digit was heterogeneous in MD cells (indicating a preferred digit), with the firing rate disparity increasing with the RF size of cells. The relationship between firing rate and cells’ RF size was also evident in the baseline activity of the preferred digit, and in the 2^nd^ preferred digit, indicating that gain properties of MD cells are generalized across the somatotopic RF. Taken together, these data suggest that response gain functions of area 3b cells are dependent on their RF size, becoming more excitable, likely, by accumulating afferent synaptic activity from connected cells with similar somatotopic RFs.

The leading view in the somatosensory system indicates that area 3b predominantly operates as a ‘local feature detector’, relaying signals to downstream areas where cross-finger neural integration emerges (Hsiao, 2008; Hsiao and Gomez-Ramirez, 2013). Our data argues against this model by showing that a majority of area 3b cells have RF that span multiple digits. The question arises: Why hasn’t this large incidence in MD RFs in area 3b been reported in the literature? We do not have a conclusive answer, but we suspect that standard practices in mapping somatosensory RFs plays a large role. Most studies describing the RF properties of area 3b have used anesthesia. Yet, anesthesia can have a significant impact on neurons’ activation strength (Hildebrandt et al., 2017; Krom et al., 2020; Sorrenti et al., 2021), and, thus, neural responses to non-preferred digit stimulation may have been substantially suppressed by the anesthetic. Another major factor is that most, if not all, studies have used unreliable hand-held probes to initially map the classical RF properties of cells (see e.g., (Ageranioti-Bélanger and Chapman, 1992; Bensmaia et al., 2008; DiCarlo et al., 1998; Pei et al., 2009, 2010, 2011; Sripati, 2006; Tabot et al., 2013)). As we have shown, this approach can lead to an inaccurate view of the RF properties of neurons in the somatosensory system (Kim et al., 2015). Using motorized devices we demonstrated that neurons in areas 3a and 3b of SI integrate information from both cutaneous and proprioceptive channels, arguing against the modality-specificity model of the somatosensory system (Mountcastle, 2005; Sur et al., 1981, 1984). Our findings are in agreement with Reed and colleagues (Reed et al., 2010a), and show that MD RFs of area 3b extend to classical RF structures, with cells’ excitation strength dependent on RF size. Our data argue in favor of using automated devices to systematically map cells’ RFs in the somatosensory system. This practice in mapping RFs will provide a detailed account of cells’ classical and non-classical RF properties, which can be used to generate more comprehensive models of haptics-related functions in the somatosensory system.

### Temporal Properties of the RF size in MD Cells

Responses to tactile stimulation on the preferred digit were faster in MD vs. SD cells, with inhibited responses emerging at an earlier time point as compared to excited responses. Timing differences between excited and inhibited activations may be driven by preferential projections from 1^st^ order thalamo-cortical (TC) neurons in the ventro-posterior (VP) nucleus of the thalamus. Although there is an approximate 4-to-1 excitatory/inhibitory cell ratio in cortex (Meinecke and Peters, 1987), studies show that 1^st^ order TC cells project more strongly and with higher prevalence to inhibitory cells (Bruno and Simons, 2002; Cruikshank et al., 2007; Gabernet et al., 2005; Porter et al., 2001). In support of this observation, TC excitatory currents in interneurons emerge fast, allowing inhibitory cells to fire spikes before feedforward inhibition emerges (Cruikshank et al., 2007). The same is not true for TC excitatory currents in pyramidal cells (Cruikshank et al., 2007). The implication of these data is that TC synapses drive interneurons much faster as compared to excitatory cells. As such, in neocortical areas with a higher number of excitatory vs. inhibitory cells (Meinecke and Peters, 1987), but where interneurons densely synapse onto local pyramidal cells (>50% within ∼100□m) (Isaacson and Scanziani, 2011), the population’s initial net response would be largely inhibited. This hypothesis nicely fits our observation that inhibited responses emerge earlier in area 3b cells. The finding that inhibited responses in MD cells onset earlier suggests that TC projecting cells are preferentially biased towards area 3b cells with a RF across multiple digits, with these TC-projecting inputs driving area 3b cells with higher reliability as compared to those of SD cells (**Figure 4B**).

A major finding of this paper is that the number of digit responsivity of MD cells (i.e., the RF size of MD cells) increased as a function of time (**Figure 5**), with responses systematically delayed across non-preferred digits. This finding suggests that MD RF are not generated by feedforward projections from TC relay cells, but rather from neural sources that provide feedback to area 3b. Although we cannot ascertain the source that provides feedback to area 3b MD cells, a recent study in mice showed a direct and somatotopically-aligned circuit between SI and SII (Minamisawa et al., 2018). This connectivity pattern has also been described in rats (Liao and Yen, 2008), with 1^st^ order TC relay cells projecting in parallel and independently to SI and SII cortices. However, a recent study in non-human primates indicates that area 1 may represent a feedback source to area 3b by relaying the integrated signals from extensive somatotopic locations (Pálfi et al., 2018). Area 3b receives inputs from both area 1 and SII, thus MD RFs in this area may arise from a combination of inputs from these two areas. Clearly, further studies are needed to identify the feedback source(s) and laminar target(s) that give rise to MD RFs in area 3b cells.

### Orientation Selectivity Differs between SD and MD area 3b cells

The data revealed that orientation tuning was more prevalent in SD as compared to MD cells (**Figure 6B)**. SD cells also had increased orientation selectivity (**Figure 6C**), suggesting that cells with a SD RF have higher spatial acuity functions. These findings are consistent with human anecdotal and experimental observations showing that fine analyses of an object feature (e.g., textures, edges, pressure) is usually performed by scanning the surface with a limited number of fingers (Lederman and Klatzky, 1987). Of note, although studies show that the distal finger pad of the index and middle fingers (i.e., D2 and D3) have the highest density of mechanoreceptors (Johansson and Vallbo, 1979), and, thus, greater sensitivity, we failed to observe higher OI in cells with a RF over D2 or D3 (**Supplemental Figure 6**).

Orientation preference across fingers was correlated in MD cells, but largely phase-shifted between 22.5° and 67.5°, with the most common difference at 22.5° **(Figure 6D)**. This cross-finger orientation angle disparity is different than the cross-finger angle preference in SII cortex (Fitzgerald et al., 2006), wherein cells’ orientation preference is highly similar across responsive finger pads (i.e., angle disparity between fingers ∼0°). These cross-finger coding discrepancies could arise neural transformations occurring between area 3b and SII. Although there are direct connections between area 3b and SII (Jones and Powell, 1969a, 1969b), there is at least one synapse between downstream outputs of area 3b to SII (e.g., area 3b → area 1 → SII). Thus, orientation selective populations in area 1 might transform the cross-finger orientation disparity of area 3b cells into a common orientation preference map that is then relayed to SII. A more provocative explanation is that area 3b cells may receive feedback input from SII neurons that encode different orientation angles. Indeed, phase-shifted orientation preference across digits can be a useful coding property for representing object shape.

We propose that cells that integrate inputs from multiple fingers, and have cross-finger orientation preferences that differ between 22.5° and 67.5° are advantageous (e.g., less computationally demanding) for representing non-squared objects (e.g., triangular or hexagonal-shaped object). For instance, a population readout that represents a non-squared object could be achieved by sampling from a limited set of MD neurons that have relatively similar orientation tuning preferences in the preferred digit but phased-shifted preferred orientations in the non-preferred digits. In contrast, neural representations of non-squared objects derived from a neural population with orientation tuning properties similar to those in SII would require readout from a larger set of cells that differ in orientation preferences, thus making the process more computationally expensive. Indeed, this latter mechanism requires a selective and convoluted combination of cells that have orientation preference disparities between 22.5° and 67.5 across each digit.

## Conclusion

The traditional view in the somatosensory system indicates that object perception emerges across cortical stages, with area 3b serving as a ‘feature detector’ cortical relay site (Hsiao, 2008). Recent studies push against this framework by showing that neurons in area 3a and 3b integrate inputs from both cutaneous and proprioceptive modalities (Kim et al., 2015). Our study further argues against this ‘serial model’ of object perception by demonstrating that area 3b has cells that encode orientation signals across multiple fingers using a coding scheme that may be amenable for representing curved objects. Further, we found higher orientation selectivity in SD vs. MD cells. Based on these observations, we propose two parallel functional pathways in area 3b: (1) A ‘*Spatial Acuity’* pathway, comprising SD cells, that is important for providing a detailed representation of particular tactile spatial features; and (2) An ‘*Object-Shape Formation*’ pathway, comprising MD cells, that is integral to the process of generating a neural representation of the 3D contour of tactile objects.

## Methods

### Animals

Experiments were conducted in two male rhesus (Macaca mulatta; 8.1 – 9.7 kg weight) monkeys. Each monkey underwent surgery under anesthesia to implant head restraining posts and recording chambers. Recording chambers (stainless steel, 19 mm diameter) were placed on both hemispheres to target the hand region of somatosensory cortex, centered over the Horsley-Clarke coordinates anterior/posterior = 6, medial/lateral = ±22. All surgical and experimental procedures were approved by the Animal Care and Use Committee of the Johns Hopkins University and conformed to National Institutes of Health and U.S. Department of Agriculture guidelines.

### Experimental design and stimuli

Animals sat in a comfortable chair with their head restrained, and tested hand in a supinated posture. The fingers on the tested hand were placed comfortably in a hand holder, with digits 2 through 5 secured with cloth tape around the medial pads, and the fingernails secured with a small amount of fixative to a nail holder. A custom-made tactile stimulator device was used to systematically map the RF and orientation tuning properties of cells on the distal pad of D2-D5 (**Figure 1)**. The tactile stimulator was a linear motor mounted on the shaft of a rotating stepper motor (Arsape AM 1020, 10 mm diameter, 15.9 mm length, Faulhaber, Clearwater, Fl.). The oriented bar was 3D printed (Objet Alaris 30U), 10mm long, approximately the width of a monkey’s finger, with its short axis equal to 3 mm. A wedge-shaped bar (90°) was used to produce a robust edge sensation of a surface. The motors were attached to an articulated tool holder (Noga Engineering Ltd. Shlomi 22832, Israel) mounted to a micro-positioner (Newport Corp., California) and a magnetic base. The animal’s fingers were slightly spread to avoid the bar contacting multiple digits. The bar was presented for 500 ms, with a ramp time of 20ms, 1mm indented into the skin, and with an orientation between 0° and 157.5° (steps of 22.5°). Each orientation was presented eight times. The interstimulus interval (ISI) was 700 ms.

Animals were trained to perform a visual discrimination task while being presented with the tactile stimuli. The visual task began with presentation of a blue square with size of 2.04°. After 400 ms of fixation, two white circles appeared on the left and right of the visual cue (each 2.04° in diameter). The circles had different luminance levels, and the animal was required to make a saccade to the brighter circle. The two visual circles were presented for a maximum of 2000 msec. The inter-trial-interval was 2300 msec. The discrimination difficulty was adapted using an ongoing staircase method based on the animal’s performance (Gomez-Ramirez et al., 2014). The difficulty increased (i.e. the luminance difference decreased, using a logarithmic scale) following three successive correct trials, and decreased after each error. The animal was rewarded with a drop of liquid after every correct response. All visual stimuli were presented on a Samsung SyncMaster 740b 17” LCD monitor, on a black background with a 60 Hz refresh rate. Eye position was monitored with a PC-60 ViewPoint EyeTracker (Arrington Research - Scottsdale, AZ).

### Neurophysiology

Neural activity was recorded from area 3b from four hemispheres (see supplementary material for details on training in each animal). Standard neurophysiological techniques were used to collect the data in all animals (Gomez-Ramirez et al., 2014; Kim et al., 2015; Ray et al., 2008; Steinmetz et al., 2000; Thakur et al., 2006). Prior to recording, a craniotomy was made in the center of the recording chamber, approximately 3 mm in diameter. Thereafter, the animal was brought in daily to the laboratory (6 days/week), the chamber cover removed, the chamber rinsed with sterile saline, and a positioning stage mounted onto the chamber. A custom-built microdrive system was secured to this positioning stage, containing four separate extracellular microelectrodes (2 to 7MΩ, Tungsten FHC Inc, Bowdoin, ME) linearly aligned and spaced 584 µm apart. The animal was transferred to the recording room, and the electrodes advanced through the intact dura and cortex. Electrodes were removed, the chamber cleaned, and a small piece of gelfoam with dexamethasone and antibiotic was placed on the dura the end of each recording day. The chamber was filled with sterile saline and sealed, and the animal placed back in its home cage. As recordings progressed, the initial craniotomy was expanded to cover the 3b hand region.

We used similar functional mapping procedures as in previous studies to ensure we recorded single units from area 3b, which resides in the postcentral gyrus (Bensmaia et al., 2008; DiCarlo et al., 1998; Pei et al., 2009, 2010, 2011; Thakur et al., 2012). The central sulcus was determined by the depth and transitions of white and grey matter, as well by the presence of motor responses in anterior electrodes. We positioned the recording array in an anterior-posterior orientation to predominantly cover neural representations from the same digit. It took approximately a week to localize the neural representation of digits 2-5 (D2-D5) of area 3b. After localizing D2-D5 of area 3b, the electrode array was shifted 100 m on each recording day, on the anterior-posterior axis, to track the postcentral gyrus until the entire 3b representation of D2-D5 was covered (see **Supplemental Figure 5**). We recorded activity from each hemisphere for about 2 months.

Electrodes were positioned to penetrate the brain perpendicularly. Electrodes contacted area 1 (or sometimes area 2), which lie in the surface of the brain. As one descends from the cortical surface through area 1, RFs progress from the distal, to middle, to proximal finger pads, and then to the palmar whorls. When electrodes reach area 3b, RFs proceed back up the finger, transitioning from proximal, to medial, and ultimately to distal pads. This reversed topographical representation between distal and proximal pads is a hallmark of the boundary region between area 3b and 1 (DiCarlo et al., 1998). Analyses were performed from cells located > 2mm below the surface of the brain (i.e., area 1) to further ensure we selectively sampled activity from area 3b. We focused our recordings on cells with a RF on the distal pads of D2-D5. Due to technical and logistical challenges, we did not perform histological analyses to anatomically confirm that recordings were made from area 3b. However, we reiterate that we used similar functional mapping procedures as previous experiments that studied response properties of area 3b cells (Bensmaia et al., 2008; Kim et al., 2015; Pei et al., 2009, 2010, 2011; Thakur et al., 2012), and the temporal and topographical properties of our neural recordings are highly consistent with the response properties of area 3b cells.

### RF mapping and orientation selectivity

We first isolated single units with a RF on the distal pads of D2 – D5 via manual mapping procedures. Afterwards, a custom-made tactile stimulator device was used to deliver tactile stimuli of different orientations to the distal pad of D2-D5. We only analyzed cells where the orientation selectivity and RF mapping procedure was fully conducted in D2-D5 distal pads.

A large fraction of somatosensory cells have inhibited responses to the presentation of a tactile stimulus (Fitzgerald et al., 2004, 2004; Gomez-Ramirez et al., 2014; Ray et al., 2008; Thakur et al., 2006). Thus, we computed the absolute value of baselined responses to capture the firing rate power elicited by the stimulus regardless of response directionality (i.e., inhibited vs. excited; equation 1).

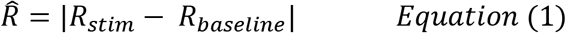

where R̂ is the absolute value of the difference between the baseline firing rate (R_baseline_) and the firing rate during the stimulus (R_stim_). We defined the cell’s ‘preferred’ digit as the stimulated digit that evoked the highest R̂. The cell was classified as ‘excited’ or ‘inhibited’ depending on whether the post-stimulus response was greater (excited) or lower (inhibited) to the mean activity in the baseline period.

We estimated a neuron’s response similarity across digits by quantifying the Fano Factor across responsive digits (see equation 2).

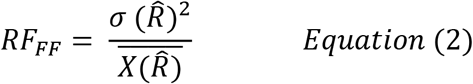

□^2^ and **X̅** represent the variance and mean across all trials in the digits that evoked a significant response in that cell. A value of one, indicates a MD cell with Poisson distributed responses across digits. A number below one indicates a MD cell with similar responses across digits, whereas a value greater than one represents a MD cell with an unequal response across digits (i.e., a MD cell with a preferred digit).

Orientation selectivity for each digit was computed using equation 3,

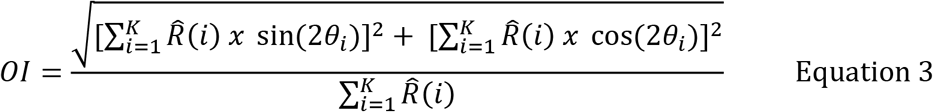

where R̂(i) is the average baselined absolute firing rate in response to the bar at orientation Θ_i_. We used the average response over the entire stimulus period (onset to 100msec after offset), as it has been shown that orientation selectivity can vary over the stimulus period (Bensmaia et al., 2008). Values of the orientation index (OI) range from 0, where a neuron has a uniform response to all orientations, to 1, where the neuron has a non-zero response to only 1 orientation. For each neuron, we determined the statistical significance of OI by randomizing responses across repetitions 5000 times and recalculating OI each time to obtain a distribution of values expected by chance (Yau et al., 2009). A separate randomization distribution was calculated for each cell. We defined tuning to be significant when the observed OI value exceeded 95% of the values in the randomized distribution. The preferred orientation was determined as the orientation that elicited the highest response. Circular correlation methods were performed to determine the similarity in orientation tuning preference between preferred and non-preferred digits of MD cells.

### Ensuring single unit (SU) isolation

We performed several methods to ensure that recordings across all sessions were obtained from the same cell (Gomez-Ramirez et al., 2014). Implementing ‘sanity check’ methods for SU identification is critical, as multi-unit recordings may lead to the incorrect assignment of a cell with a MD RF. SUs were isolated using a template-based spike sorter, and only one neuron per electrode was recorded at a time (Thakur et al., 2007). The shape and timing information of each action potential (AP) was stored, and additional SU isolation analyses were performed offline to ensure that SU activity was well isolated. First, spikes occurring within 3msec of one another were excluded, as it would be unlikely to observe such spikes from the same neuron. Next, the shape of the AP was subjected to principal component analysis (PCA), and shapes that were more than three standard deviations away from the center of mass of the two most principal components (using the normalized Euclidean distance method) were deleted. Next, the experimenter visually inspected each block of trials, and manually deleted AP shapes that were deemed outliers. Finally, we sorted the mean firing rate of a cell across trials within one protocol, and fitted the sorted firing rates with power function. Trials from the tail ends were deleted until the fit produced a non-significant fit (p > 0.05). Since the experimental conditions were uniformly randomized, a negative or positive slope of the sorted trials would be indicative of cell loss or inclusion of APs from nearby cells, respectively. Cells with less than 30 trials were not included in the analyses. Our analyses led to 477 SUs from the hand regions of area 3b, of which 223 neurons were recorded from two hemispheres in animal MR4358M, and 254 were recoded from two hemispheres in animal 43V.

### Determining the temporal evolution of neural responses

We performed a temporal clustering analysis to determine whether MD responses emerge across time vs. simultaneously. On each trial, we epoched the data between stimulus onset to 160ms after stimulus onset (to ensure that ‘Off’ responses were included in the analyses), and divided the epoch into 20ms bins. For each bin, we determined whether the average response was significantly different than baseline (using Wilcoxon rank-sum test, p <0.05). The baseline activity was computed as the average response between -450 and -250msec, relative to stimulus onset. If a cell had two consecutive bins with a significant response in the same direction with respect to baseline (two consecutive negative or positive bins), we considered that cell to have a significant response on that digit (Kim et al., 2015).

## Supplemental Material

### Results

#### Example of cells with different RF size

**Supplementary Figure 1** shows additional example cells with single digit (SD) and multi-digit (MD) RFs. This figure shows two examples cells for each RF size category. The top row graphs show cells with a SD RF (Left graph: Cell with a RF over D4; Right graph: Cell with a RF over D2). The 2^nd^ row from the top shows cells with a RF spanning two digits (Left graph: Cell with a RF over D3 and D4; Right graph: Cell with a RF over D2 and D3). The 3^rd^ row from the top shows cells with a RF spanning three digits (Left graph: Cell with a RF over D2, D4, and D5; Right graph: Cell with a RF over D3, D4 and D5). The bottom row graphs show cells with a RF spanning four digits (Left graph: Cell with a RF over D2, D3, D4, and D5; Right graph: Cell with a RF over D2, D3, D4, and D5). The insets in each graph depict the averaged action potential waveforms recorded to each responsive digit.

**Supplemental Figure 1:**
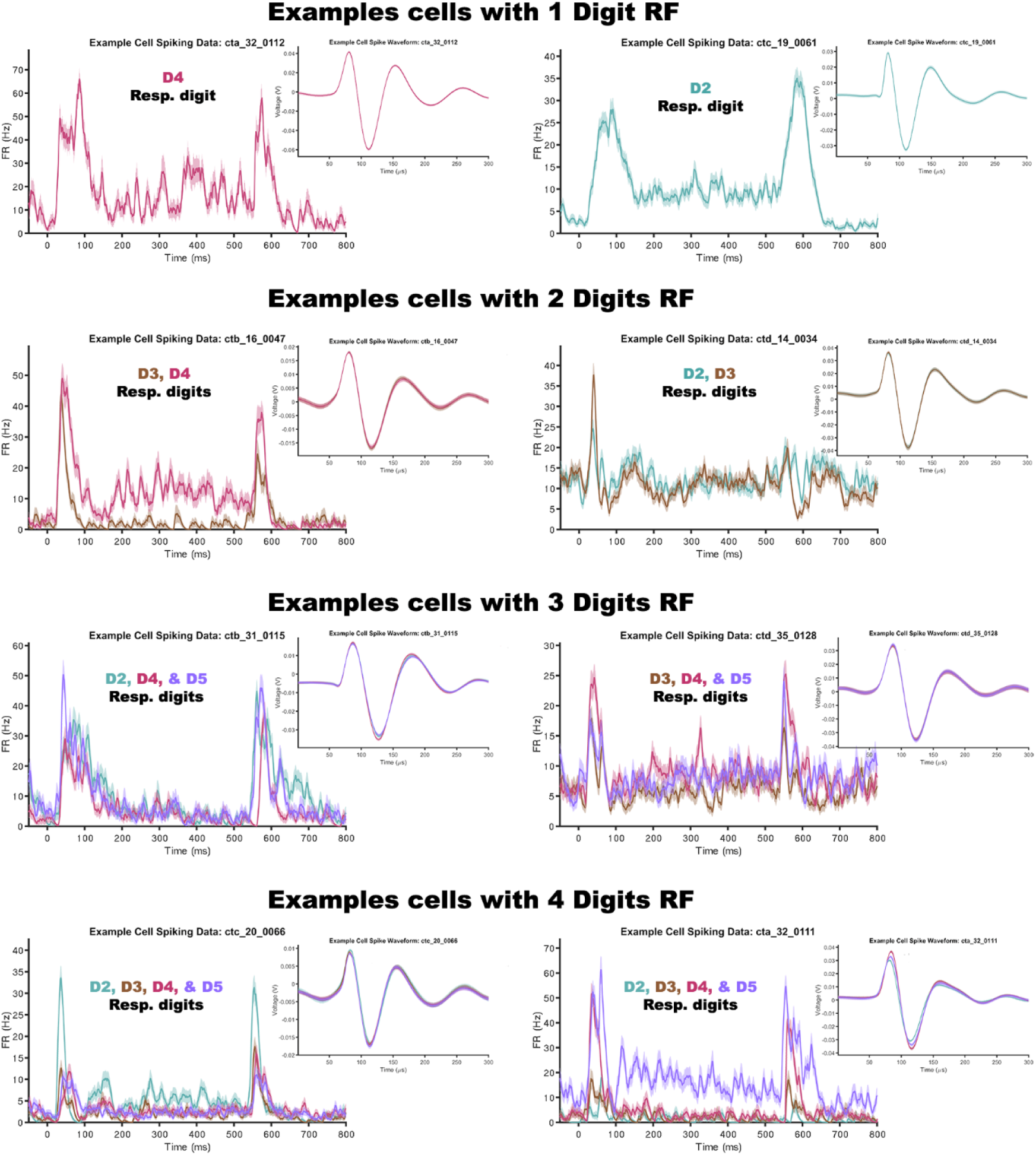

#### RF Size Distribution of Cells is similar across all Hemispheres

The pattern of RF size distribution was similar across all hemispheres and monkeys. **Supplementary Figure 2**, shows the percentage of cell with one, two, three, and four digits RF for Monkey MR4358M (top graphs), and Monkey 43V (bottom graphs). The right-most bar in each of the graphs represents the percentage of cells with a RF spanning 2 or more digits. We recorded activity in two of the hemispheres (CXA and CXD) in an animal that was trained in a task with stimuli delivered to multiple fingers. A similar type of tactile stimulation has been shown, in a different non-human primate species, to promote expansion of RFs (Jenkins et al., 1990; Wang et al., 1995). As **Supplementary Figure 2** shows that MD effects were not driven by ‘training’ confounds. Cells with MD RFs were clearly present in all hemispheres.

**Supplemental Figure 2:**
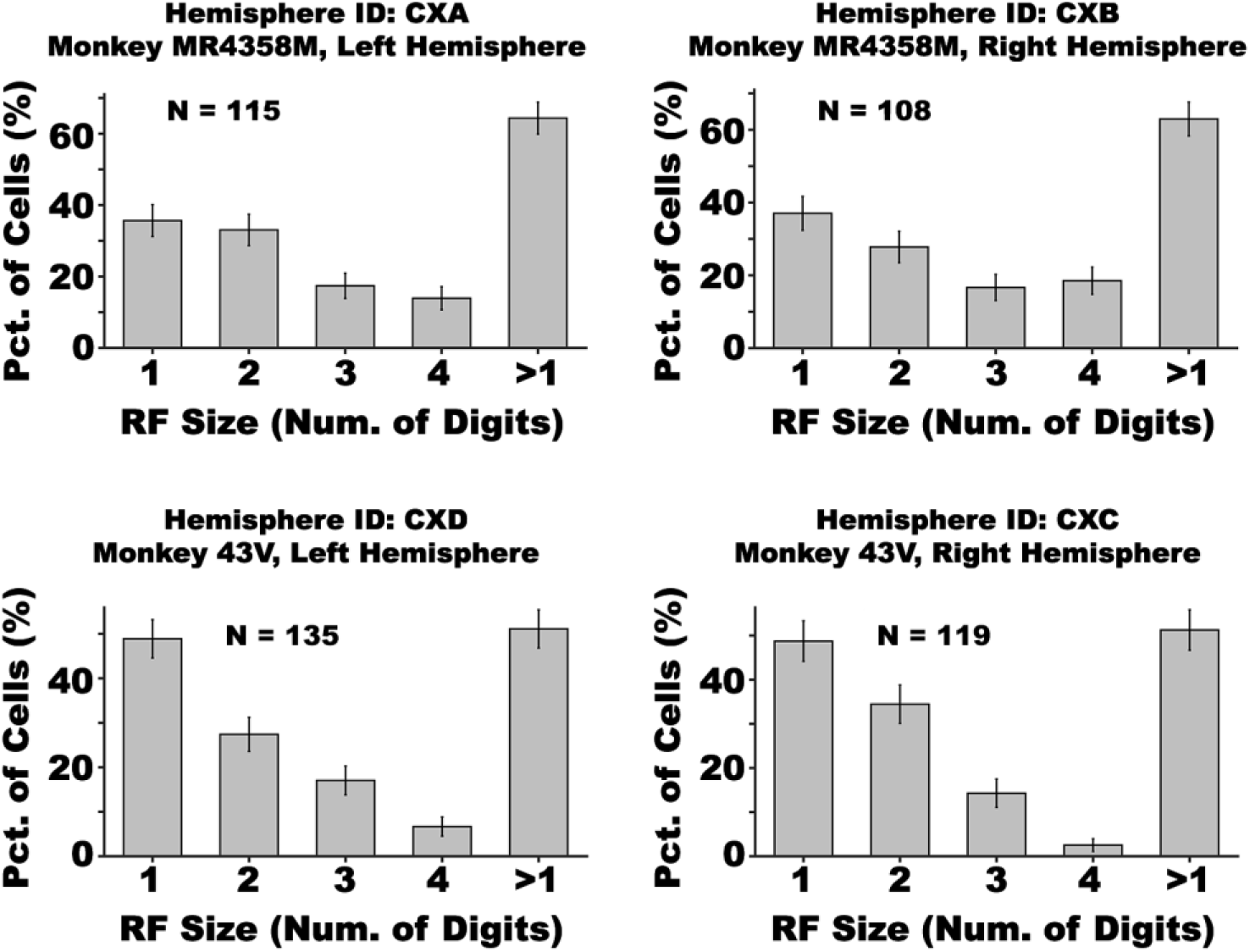

#### Spontaneous Activity during the Baseline Period increases as a Function of RF Size

The baseline firing rate was significantly different between RF size of cells (H(3, 476) = 22.92, p < 0.001). **Supplementary Figure 3** shows violin plots of the baseline firing rate in cells with a RF spanning 1, 2, 3, and 4 digits. Pairwise statistics showed differences in firing rates between cells with a RF spanning 4 vs. 2 digits (Z = 3.11, p < 0.01), 4 digits vs. 1 digit (Z = 4.06, p < 0.001), 3 vs. 2 digits (Z = 2.00, p < 0.05), and 3 digits vs. 1 digit (Z = 3.31, p < 0.001).

**Supplemental Figure 3:**
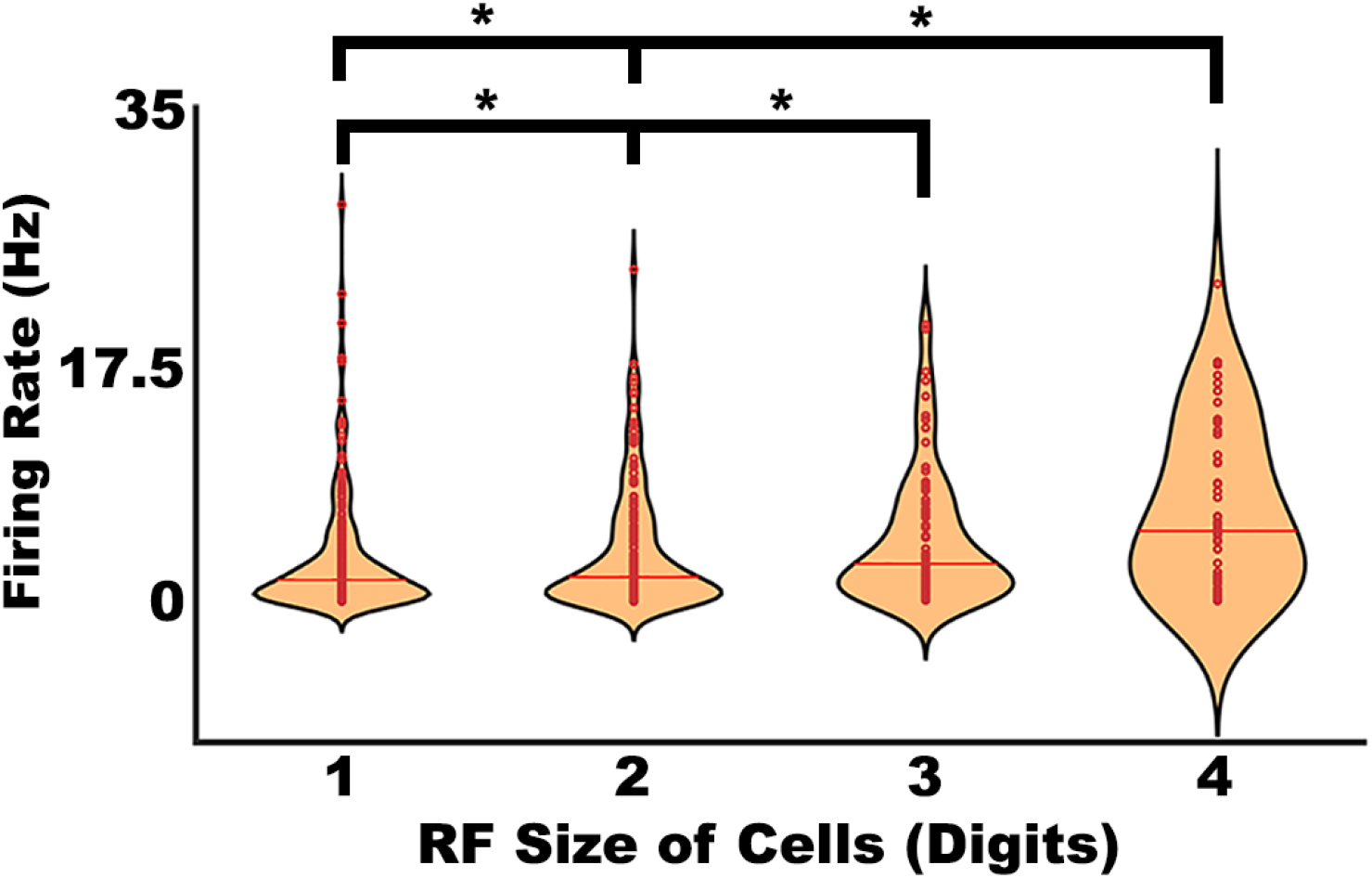

#### Neural Responses to Tactile Stimulation on the 2^nd^ Preferred Digit Increase with RF Size

Neural responses to stimulating the 2^nd^ preferred digit increased as a function of RF size (H(2,271) = 46.03, p < 0.001). **Supplementary Figure 4** shows violin plots of the normalized firing rate responses to the 2^nd^ preferred digit in cells with a RF spanning 2, 3, and 4 digits. Pairwise statistics showed differences in firing rates between cells with a RF spanning 4 vs. 3 digits (Z = 4.07, p < 0.001), 4 vs. 2 digits (Z = 6.59, p < 0.001), and 3 vs. 2 digits (Z = 2.98, p < 0.01).

**Supplemental Figure 4:**
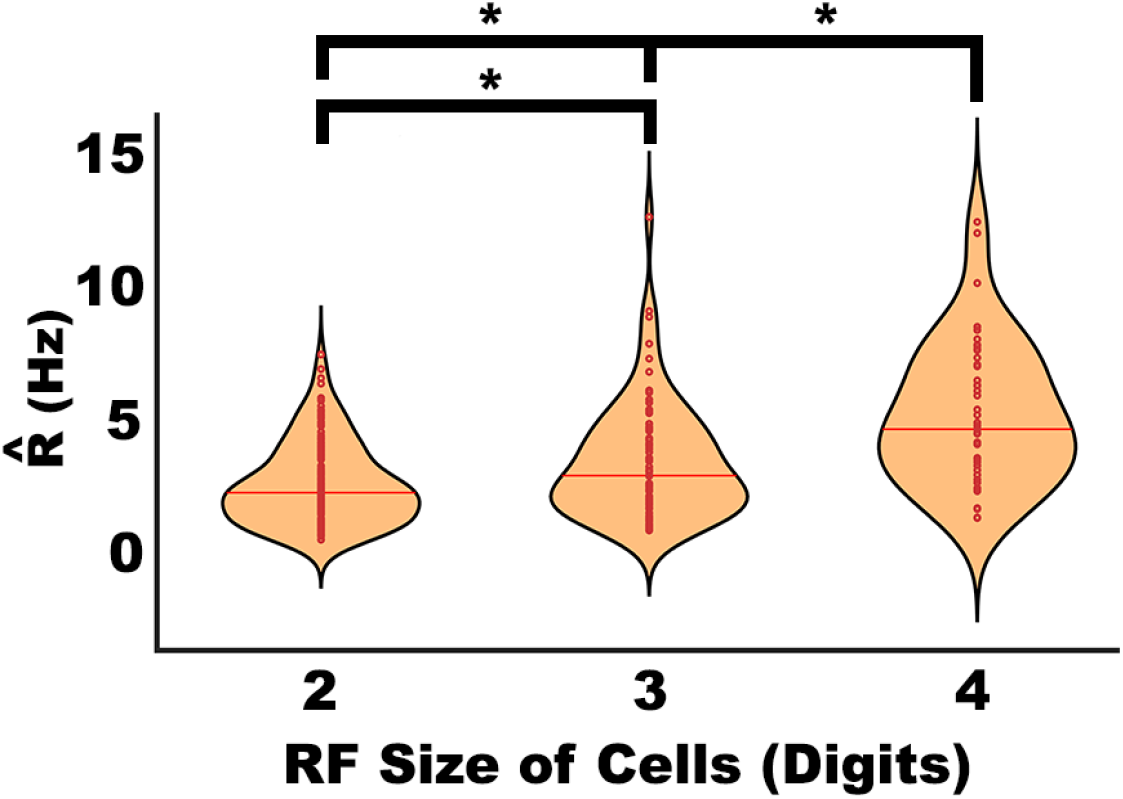

#### Each digit representation was homogenously sampled across the finger area of 3b

**Supplementary Figure 5A** shows the percentage of cells with the preferred digit location over D2, D3, D4, or D5. No Differences were observed between the percentages of cells with a preferred digit in a particular location. **Supplementary Figure 5B** shows the percentage of cells with the digit location of the 2^nd^ most responsive digit relative to the location of the preferred digit. The negative and positive values indicate whether the 2^nd^ preferred digit is to the left or right of the preferred digit, respectively.

**Supplemental Figure 5:**
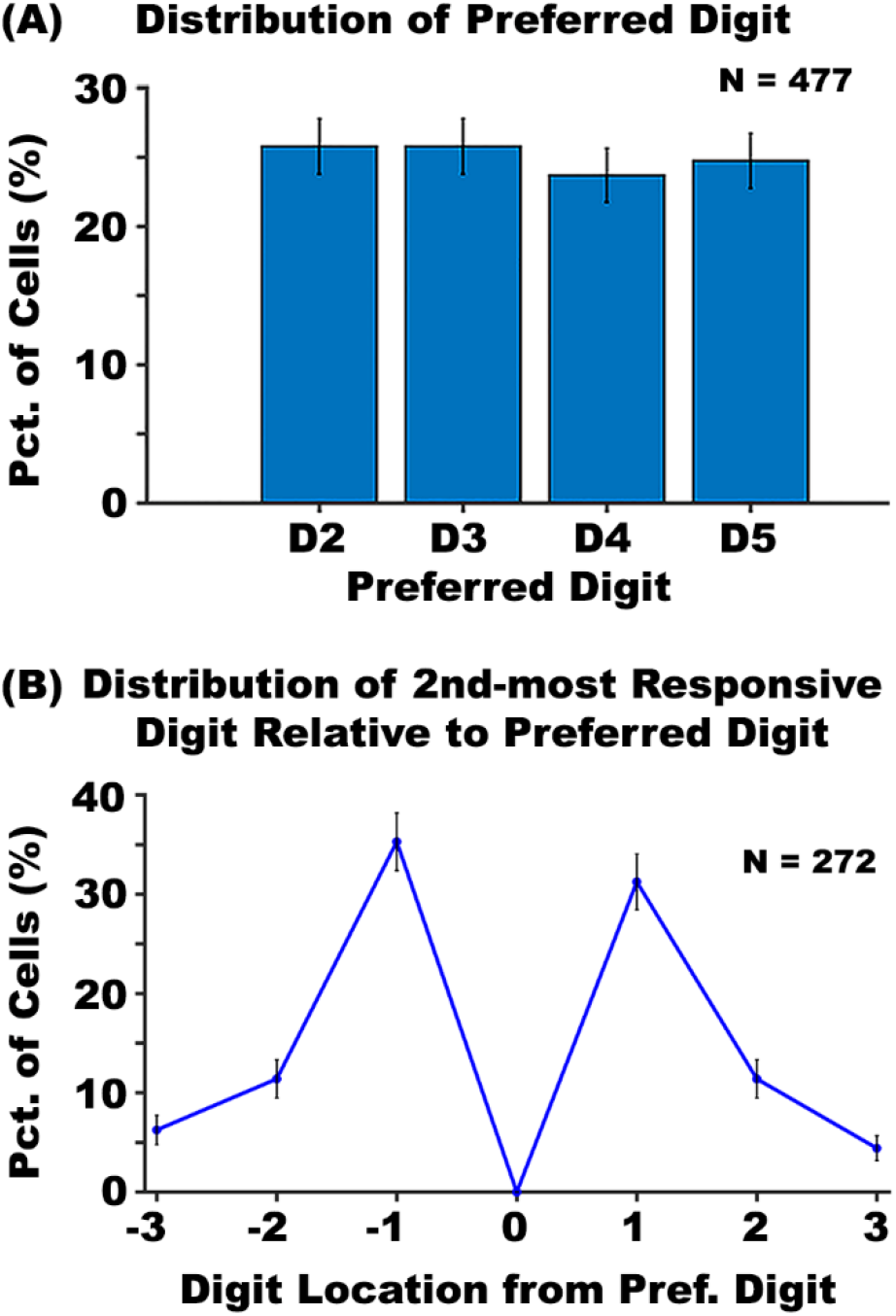

#### Orientation Selectivity is Similar across All Digits

**Supplementary Figure 6** shows the Orientation Index (OI) for each digit collapsed across cells with a SD or a MD RF. The data did not reveal significant differences in OI between digits (H(3,458) = 2.83, p > 0.05)). This null effect was observed in SD and MD cells as well. The OI values in D2, D3, D4, and D5 were 0.242, 0.245, .232, and .220, respectively.

**Supplemental Figure 6:**
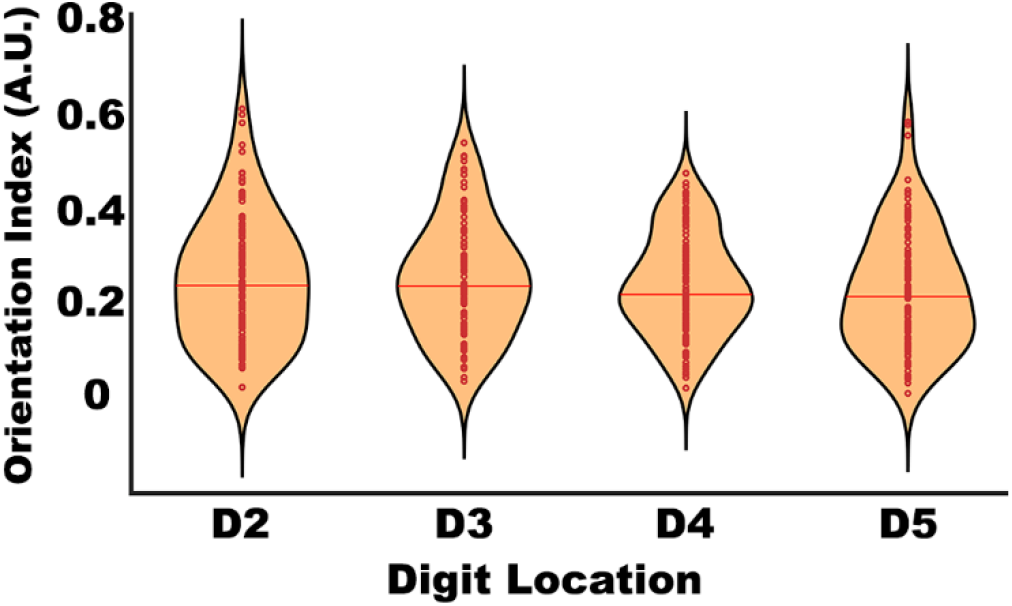

